# Predicting clinical outcomes from large scale cancer genomic profiles with deep survival models

**DOI:** 10.1101/131367

**Authors:** Safoora Yousefi, Fatemeh Amrollahi, Mohamed Amgad, Coco Dong, Joshua E. Lewis, Congzheng Song, David A Gutman, Sameer H. Halani, Jose Enrique Velazquez Vega, Daniel J Brat, Lee AD Cooper

## Abstract

Translating the vast data generated by genomic platforms into accurate predictions of clinical outcomes is a fundamental challenge in genomic medicine. Many prediction methods face limitations in learning from the high-dimensional profiles generated by these platforms, and rely on experts to hand-select a small number of features for training prediction models. In this paper, we demonstrate how deep learning and Bayesian optimization methods that have been remarkably successful in general high-dimensional prediction tasks can be adapted to the problem of predicting cancer outcomes. We perform an extensive comparison of Bayesian optimized deep survival models and other state of the art machine learning methods for survival analysis, and describe a framework for interpreting deep survival models using a risk backpropagation technique. Finally, we illustrate that deep survival models can successfully transfer information across diseases to improve prognostic accuracy. We provide an open-source software implementation of this framework called *SurvivalNet* that enables automatic training, evaluation and interpretation of deep survival models.

## INTRODUCTION

Advanced molecular platforms can generate rich descriptions of the genetic, transcriptional, epigenetic and proteomic profiles of cancer specimens, and data from these platforms are increasingly utilized to guide clinical decision-making. Although contemporary platforms like sequencing can provide thousands to millions of features describing the molecular states of neoplastic cells, only a small number of these features have established clinical significance and are used in prognostication (1-4). Extracting reliable and accurate predictions of clinical outcomes from high-dimensional molecular data remains a major challenge in realizing the potential of precision genomic medicine.

Traditional Cox proportional hazards models require enormous cohorts for training models on high-dimensional datasets containing large numbers of features. Consequently, a small set of features is selected in a subjective process that is prone to bias and limited by imperfect understanding of disease biology. High-dimensional learning problems are common in the machine-learning community, and many machine-learning approaches have been adapted to predicting survival or time to progression (5). Regularization methods for Cox models like elastic net have been developed to perform objective and data-driven feature selection with time-to-event data (6). Random forests are reputed to resist overfitting in high-dimensional prediction problems, and have been adapted to survival modeling (7). Neural network based approaches have been used in low-dimensional survival prediction problems (8), but subsequent evaluation of these methods found no performance improvement over ordinary Cox regression (9). The difficulty of deconstructing these black-box models to gain insights into disease progression or biology remains a key challenge in their adoption.

Advances in neural networks broadly described as *deep learning* have shattered performance benchmarks in general machine-learning tasks, enabled by improvements in methodology, computing hardware, and datasets (10). These networks are composed of densely interconnected layers that sequentially transform the inputs into more predictive features through adaptive learning of the interconnection parameters (see Figure 1). Deep networks composed of many layers perform *feature-learning* on high dimensional datasets to extract latent explanatory features (11), and have been successfully applied to biomedical problems including image classification (12), transcription factor binding site prediction (13), and medication dosing control (14). A fundamental challenge in deep learning is determining the network design that provides the best prediction accuracy, a process that involves choosing network *hyperparameters* including the number of layers, transformation types, and training parameters. Searching the vast space of network designs quickly becomes intractable, given the considerable time required to train a single deep network. *Bayesian optimization* techniques have been developed to automate the search of the hyperparameter space, and provide measurable performance gains over hand-tuning procedures (15). Advanced deep learning techniques including dropout regularization, unsupervised pre-training, and Bayesian optimization were first applied to build unbiased deep models from high-dimensional genomic data in (16) where deep networks were trained to optimize proportional hazards likelihood. A subsequent study applied deep networks to model survival in breast cancer using a low-dimensional dataset (14 features) that were selected with a priori disease knowledge (17). This study did not evaluate prediction using high-dimensional data or compare to state-of-the-art methods like regularized Cox regression that perform unbiased feature selection.

**Figure 1.**
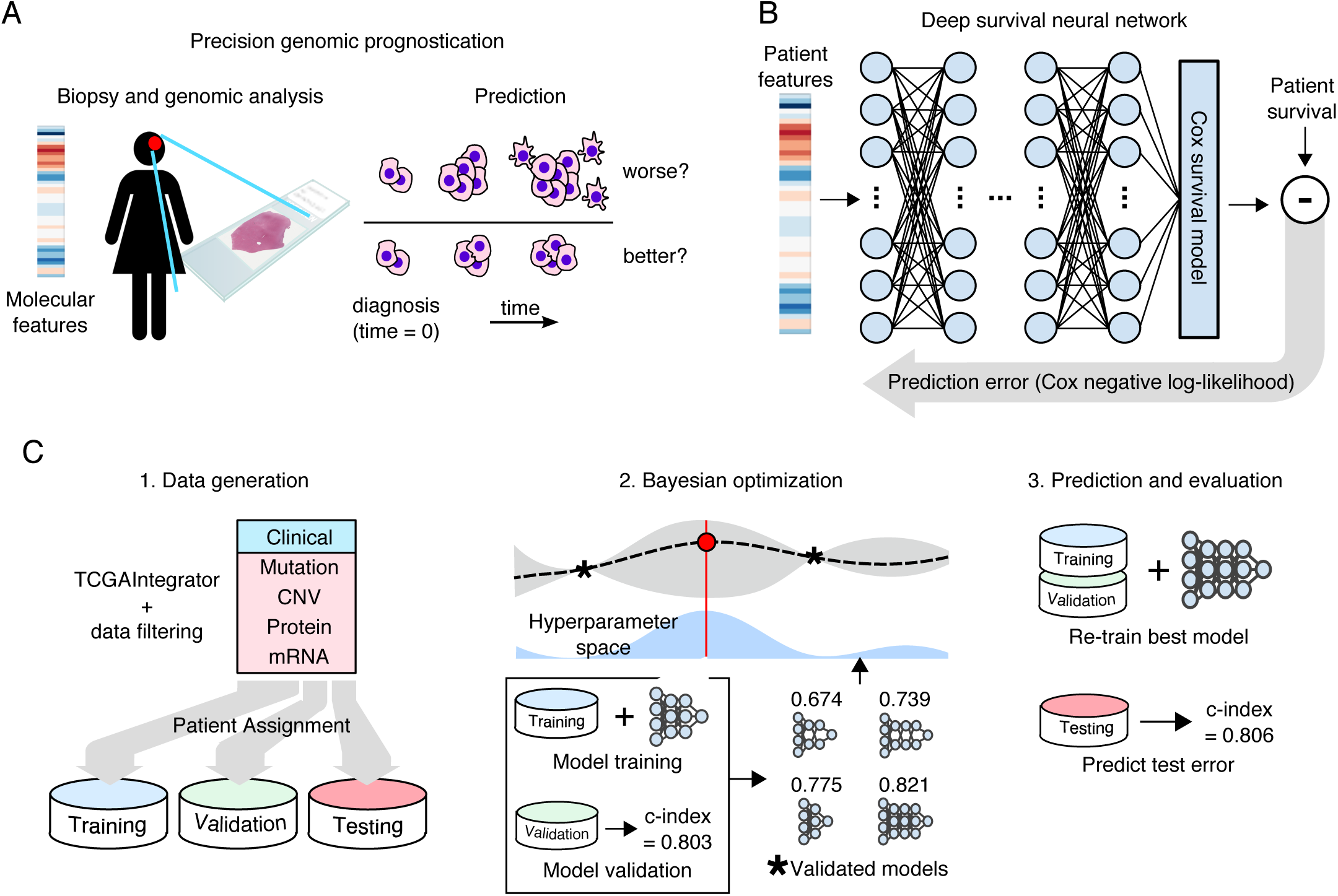
Overview of the SurvivalNet framework. **(A)** Accurate prognostication is crucial to clinical decision making in cancer treatment. Molecular platforms produce data that can be used for precision prognostication with learning algorithms. **(B)** Deep survival models are neural networks composed of layers of non-linear transformations, driven by a Cox survival model at the output layer. Model likelihood is used to adaptively train the network to improve the statistical likelihood of the overall survival prediction. **(C)** The SurvivalNet framework enables automatic design optimization and validation of deep survival models. Molecular profiles obtained from TCGA datasets are randomized, assigning patients to training, testing and validation sets. Bayesian optimization searches the space of hyperparameters like the number of network layers to optimize the model design. Each selected design is trained and evaluated using validation samples to update the Bayesian optimizer. The best model design is then evaluated on the independent testing set to measure the final optimized model accuracy.

This paper extends the preliminary studies exploring deep learning for survival modeling, and presents a software package called *SurvivalNet* (SN) that enables users to train and interpret deep survival models. SurvivalNet uses Bayesian Optimization to identify optimal hyperparameter settings, saving users considerable time and effort in choosing model parameters. We also illustrate how backpropagation methods can be modified to interpret deep survival models, scoring individual features for their contribution to risk, and show how feature risk scores can be used with pathway analysis tools to uncover higher-order biological themes associated with patient survival. Using clinical and molecular data from The Cancer Genome Atlas (TCGA), we show that Bayesian-optimized deep survival models provide comparable performance to Cox elastic net regression, and superior performance to random survival forests when analyzing high-dimensional genomic data. Finally, we show how deep survival models can learn prognostic information from multi-cancer datasets to improve prognostication through transfer learning.

## RESULTS

### Automatic training and validation of deep survival models

An overview of the SurvivalNet framework is presented in Figure 1. SurvivalNet is implemented as an open-source Python module (https://github.com/CancerDataScience/SurvivalNet) using Theano and is available as a pre-built Docker software container. A deep survival model uses the Cox partial log likelihood to train the weights of neural network to transform molecular features into explanatory factors that explain survival. The partial log likelihood serves as a feedback signal to train the model weights using backpropagation. Deep neural networks have many hyperparameters that impact prediction accuracy including the number of layers, number and type of transforming functions in each layer, and choices for optimization/regularization procedures. The time needed to train a deep survival model prohibits exhaustive hyperparameter search, and so SurvivalNet employs a Bayesian optimization strategy to identify hyperparameters that optimize prediction accuracy (18). Data is first split into training (60%), validation (%20), and testing (%20) sets. Training samples are used to train the model weights with backpropagation using the network design suggested by Bayesian optimization. The prediction accuracy of the trained deep survival model is then estimated using the validation samples, and is used to maintain a probabilistic model of performance as a function of hyperparamters. Based on the probabilistic model, the design with the best expected accuracy is inferred as the next design to test. After the Bayesian optimization process is finished (typically after a prescribed number of experiments), the best network design is used to re-train a deep survival model using the training+validation samples, and the accuracy of this best model is reported using the held-out testing samples.

### Comparing deep survival networks with Cox elastic net and random survival forests

We compared the performance of SurvivalNet models with cox elastic net (CEN) and random survival forest (RSF) models using data from multiple TCGA projects: pan-glioma (LGG/GBM), breast (BRCA), pan-kidney (KIPAN) which consists of chromophobe, clear cell, and papillary carcinomas. Datasets were selected based on the availability of molecular and clinical data and for extent of complete clinical follow up. Performance was evaluated with two feature-sets: 1) a “transcriptional” feature set containing 17,000+ gene expression features obtained by RNA-sequencing, and 2) an “integrated” feature set containing 3-400 features describing clinical features, mutations, gene and chromosome arm-level copy number variations, and protein expression features. Details of these datasets are presented in Methods and Tables S1 and S2. Optimization procedures for CEN and RSF hyperparameters are described in Methods.

In each experiment, samples were randomized to training (60%), validation (20%), and testing (20%) sets, and the performance of optimized SN, CEN, and RSF models was assessed. Performance was calculated using Harrell’s c-index, a non-parametric statistic that measures concordance between predicted risks and actual survival (19). A c-index of 1 indicates perfect concordance, and a c-index of 0.5 corresponds to random chance. Experiments were repeated for 20 randomizations to account for variations due to sample assignment. Differences in performance between methods were evaluated through rank-sum statistical testing of c-index values. Results are presented in Figure 2 (extended results are presented in Table S3).

**Figure 2.**
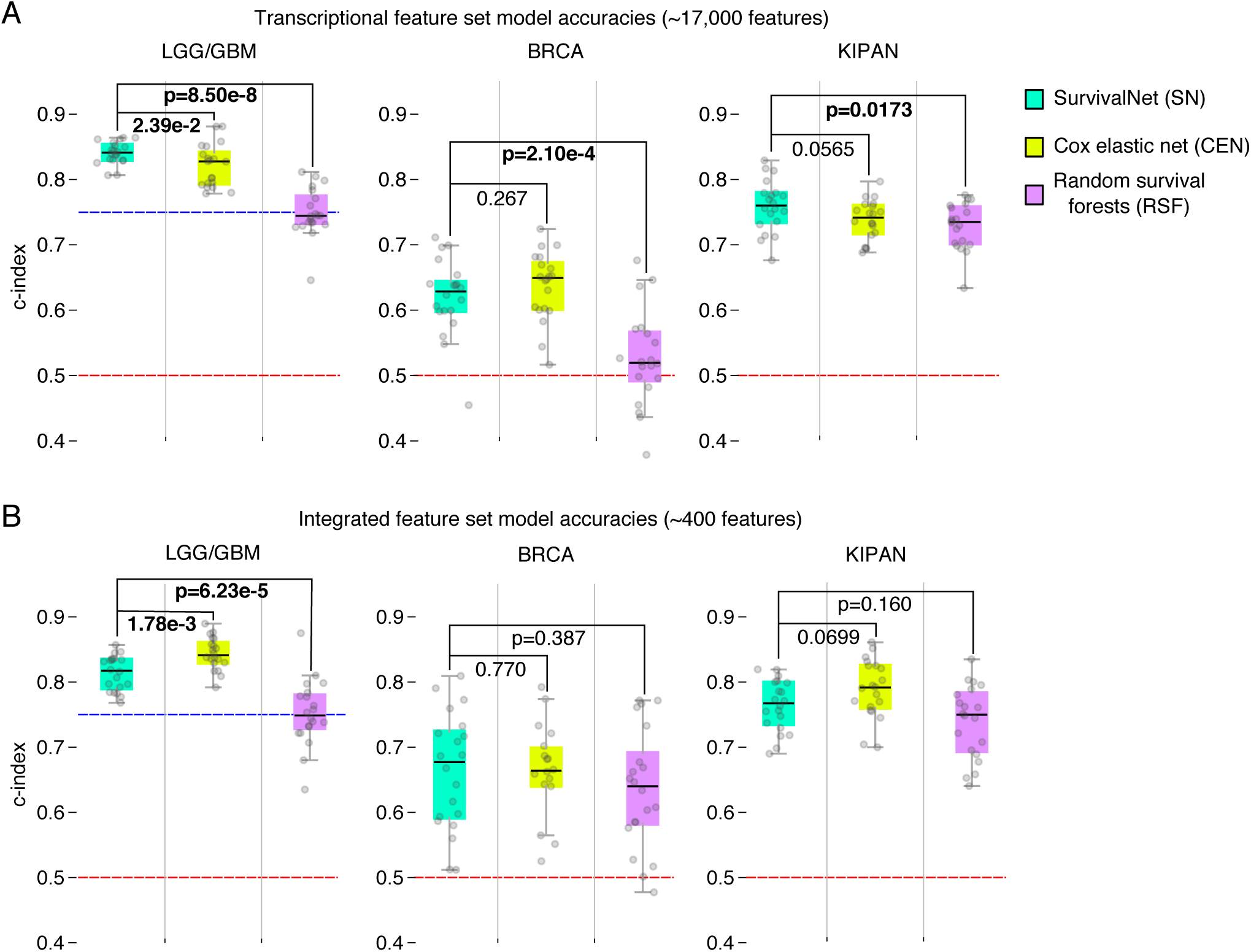
Performance comparison of SurvivalNet, Cox elastic net, and random survival forest models. The prognostic accuracy of these methods was evaluated in different diseases/datasets (GBMLGG, BRCA, KIPAN) using a high-dimensional transcriptional feature set and a lower-dimensional integrated feature set that combines clinical, genetic, and protein expression features. Patients were randomized to 20 training/validation/testing sets that were used to train, optimize, and evaluate models in each case. **(A)** SurvivalNet models have an advantage over Cox elastic net in predicting survival using high-dimensional transcriptional features. **(B)** Cox elastic net has an advantage in predicting survival using lower-dimensional integrated features. Dashed red lines corresponding to a random prediction (c-index=0.5). Dashed blue lines corresponds to c-index of molecular classification of gliomas.

Both SN and CEN significantly outperform RSF models in most experiments. All methods perform markedly better than random, with median c-index scores ranging from: 0.75-0.84 in LGG/GBM; 0.52-0.68 in BRCA; and 0.73-0.79 in KIPAN. In the transcriptional feature set (Figure 2B), SN models have a slight advantage over CEN models in LGG/GBM (**p=2.39e-2**) and KIPAN (**p=0.0565**). In the integrated feature set (Figure 2A), SN and CEN performance were indistinguishable in the BRCA dataset (**p=0.770**), but CEN models have a slight advantage over SN models in the LGG/GBM (Wilcoxon rank-sum **p=1.78e-3**) and KIPAN (**p=0.0699**) datasets. Performance is generally better on the integrated feature set than the transcriptional feature set for all methods. One exception to this is the performance of SN on the LGG/GBM feature sets, where performance on the transcriptional feature set exceeds the integrated feature set (c-index 0.841 versus 0.818). RSF models have the worst performance generally, and are severely challenged in learning from the BRCA transcriptional feature set, with a median c-index of 0.520 (slightly better than random guess).

Comparing performance across diseases, we noticed that prediction accuracy generally decreases as the proportion of right-censored samples in a dataset increases. This pattern holds for all prediction methods. Glioma had the highest overall prediction accuracy, being a uniformly fatal disease that has relatively fewer long-term survivors and incomplete follow-up (62-64%). Breast carcinoma had the lowest overall prediction accuracy with more than 86-91% of BRCA samples being right-censored.

Finally, we observed that CEN model execution routinely fails with some randomizations, producing a segmentation fault software error. In these instances, we generated a new randomization for CEN and repeated the experiments. The performance accuracy of SN and RSF models on these failed randomizations does not suggest that they present particularly difficult learning problems, but we cannot exclude the possibility of introducing a performance bias for CEN by generating new randomizations when CEN execution fails.

### Interpreting deep survival models with risk backpropagation

Linear survival models weight individual features based on their contribution to overall risk, providing a clear interpretation of the prognostic significance of individual features, and insights into the biology of disease progression. The complex transformations that machine-learning methods apply to input features makes interpreting these models more difficult. This is especially true for deep learning where the input features are subjected to multiple sequential nonlinear transformations. To enable interpretation of deep survival models, we implemented a technique that we describe as *risk backpropagation*. In the same way that backpropagation can propagate prediction errors back through the layers of a deep model for training, backpropagation can also propagate predicted risks back to the input layer to assess how individual features contribute to risk (see Figure 3). Using partial derivatives to analyze variable importance was first used in (20). In a linear model the weights of features are identical for all patients, and the risk can be conceptualized as a plane that has a uniform gradient for any input feature values. In a nonlinear model, the prediction can be conceptualized as a nonlinear surface where the risk gradients change depending on a patient’s feature values, and so these risk contributions are calculated separately for each patient.

**Figure 3.**
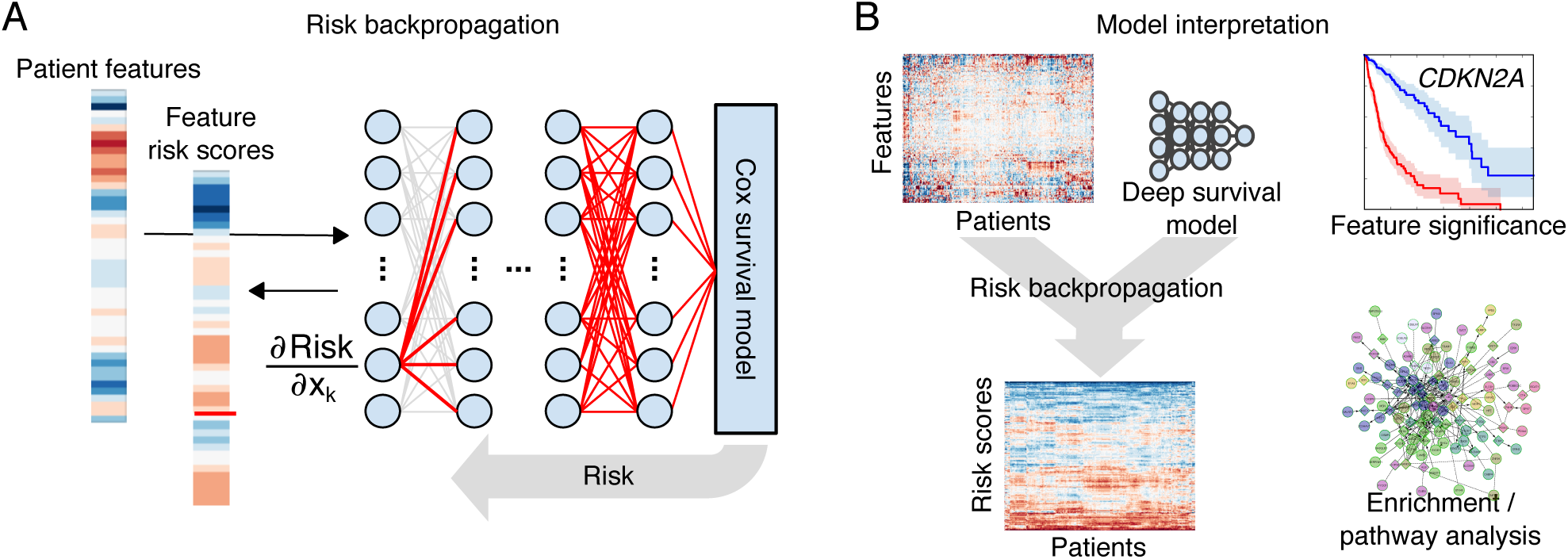
Interpreting deep survival models with risk backpropagation. **(A)** Backpropagation was used to calculate the sensitivity of predicted risk to each input feature, generating feature risk scores for each feature and patient. **(B)** Feature risk scores can be analyzed to gain insights into the deep survival model. Risk scores can be used to evaluate the prognostic significance of individual features, or to identify gene sets or molecular pathways that are enriched with high-risk or low-risk features.

We applied risk backpropagation to our integrated glioma model to investigate the prognostic significance of features in gliomas (see Figure 4). Risk backpropagation was applied to each patient to generate feature risk scores, and then features were ranked using their median score across patients as a measure of overall prognostic significance (see Figure 4A). Among the top-ranked features indicative of poor prognosis are: increased age at diagnosis (rank 3); histologic classification as de novo grade IV glioblastoma (rank 5); loss of chromosome arms 10p and 10q (ranks 2, 4); and deletions of tumor suppressor genes *CDKN2A* and *PTEN* (ranks 1, 8). The top-ranked features associated with better prognosis included mutations in *SMARCA4* (rank 6), *IDH1* / *IDH2* (ranks 9, 10) and in *CIC* (rank 17). We note that many of these features are incorporated or highly correlated with established WHO genomic classifications of gliomas (21).

**Figure 4.**
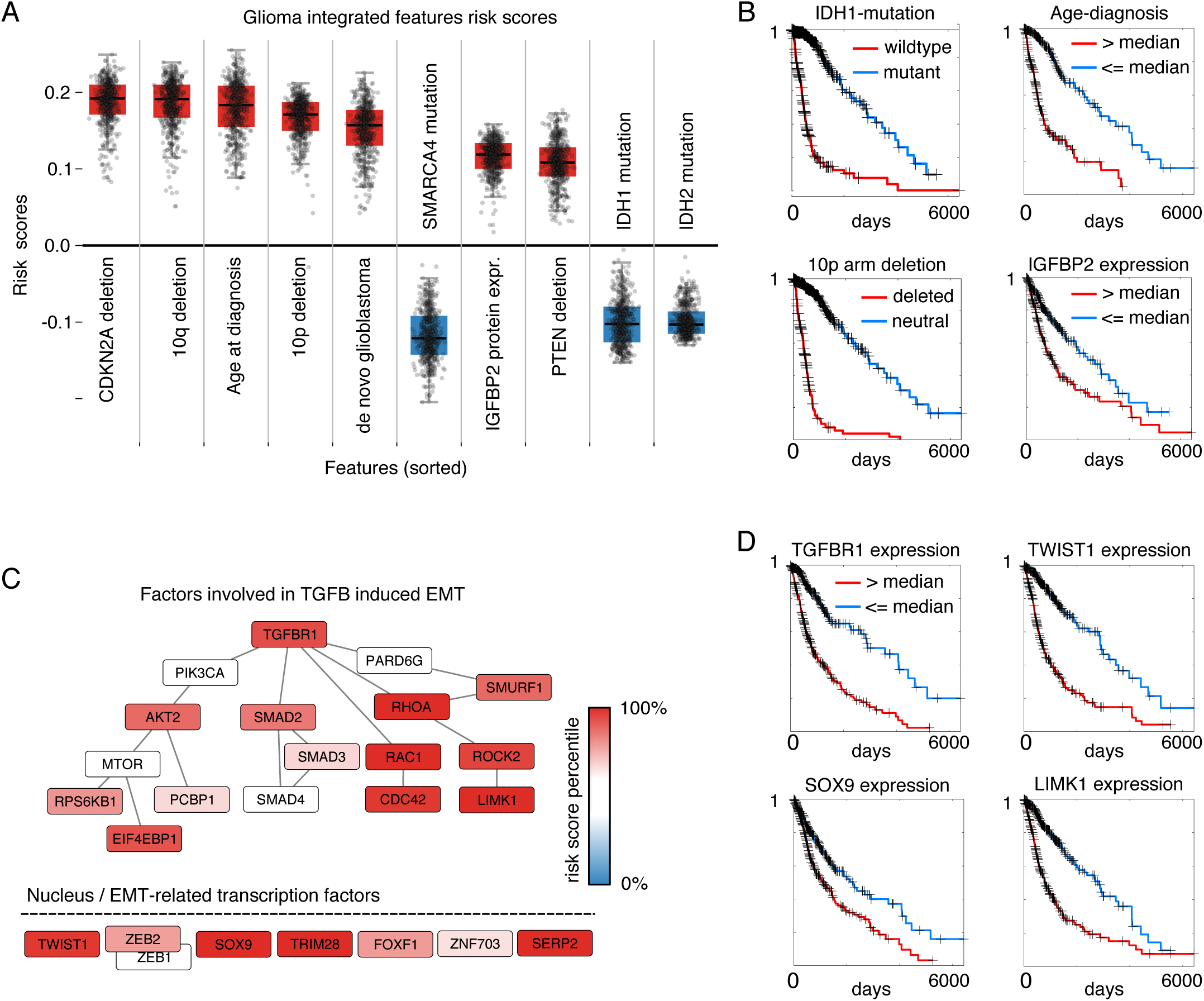
Interpretation of glioma deep survival models. (A) SurvivalNet learns features that are definitional (IDH mutation) or strongly associated (CDKN2A deletion, SMARCA4 mutation) with WHO genomic classification of diffuse gliomas. Feature risk scores for the top 10 of 399 features in the integrated model are shown here, in order. Each boxplot represents the risk scores for one feature across all patients. Features were ranked by median absolute risk score. (B) Kaplan-Meier plots for select features from (A). (C) A gene set enrichment analysis of transcriptional feature risk scores identified the TGF-Beta 1 signaling and epithelial-mesenchymal transition (EMT) gene sets as enriched with features associated with poor prognosis. (D) Kaplan-Meier plots for select features from (C).

To investigate molecular pathways related to glioma prognosis, we also performed a risk backpropagation gene-set enrichment analysis of our glioma transcriptional model. Median risk scores from the transcriptional model were calculated for each transcript, and gene set enrichment analysis was performed on these scores to identify pathways enriched with prognosis-associated transcripts (22) (See Supplementary Table 4). Pathways and gene sets associated with poor-prognosis include cell cycle (G2M checkpoint, E2F targets), apoptosis, angiogenesis, inflammation (Interferon alpha, gamma responses) and epithelial to mesenchymal transition (EMT and TGF-Beta signaling). EMT has received significant attention in cancer (23), and also specifically in glioma (24-26) as being associated with aggressive phenotypes and poor clinical outcomes. The TGF-Beta signaling hallmark gene set was significantly enriched (**p=6e-3**, FDR **q=2.7e-2**) with genes having high risk scores including *RHOA*, *TGFB1*, *TGFBR1*, *SERPINE1*, *JUNB1* and *ARID4B*. The Epithelial to Mesenchymal Transition gene set was also significantly enriched (**p=1.6e-1**, FDR **q=1.55e-1**) with genes having high risk scores including *MMP1/2/3*, *IL6*, *ECM1*, and *VCAM1*. TGF-Beta signaling is understood to be one of the main pathways involved in EMT, and our results support the importance of EMT in determining glioma patient outcomes. The feature risk scores of the EMT-related transcription factors TGF-Beta induced EMT signaling as described in (23) are visualized in Figure 4C. Major TGF-Beta-EMT inducing factors (*RHOA*, *RAC1*, *ROCK2*, *CDC43* and *LIMK1*) and EMT transcription factors (*TWIST1*, *SOX9*, *TRIM28* and *SERP2*) have among the highest risk scores in our glioma transcriptional model.

Extended feature risk scores for the LGG/GBM integrated and transcriptional models are presented in Table S4. The procedure for obtaining models used for interpretation is described in Methods.

### Transfer learning with multi-cancer datasets

We performed a series of *transfer learning* experiments to evaluate the ability of deep survival models to benefit from training with data from multiple cancer types. The transfer learning paradigm is illustrated in in Figure 5A. Survival models were trained using three different datasets: BRCA-only, BRCA+OV (ovarian serous carcinoma), and BRCA+OV+UCEC (corpus endometrial carcinoma), and were evaluated for their accuracy in predicting BRCA outcomes. The large proportion of right-censored cases in the BRCA dataset (90%) makes training accurate models difficult, and so we hypothesized that augmenting BRCA training data with samples from other hormone-driven cancers could improve BRCA prognostication. BRCA samples were randomized to training, validation, and testing and full Bayesian optimization was performed to measure c-index on BRCA testing samples for 20 randomizations. For the integrated feature set, we combined datasets by discarding disease-specific clinical features.

**Figure 5.**
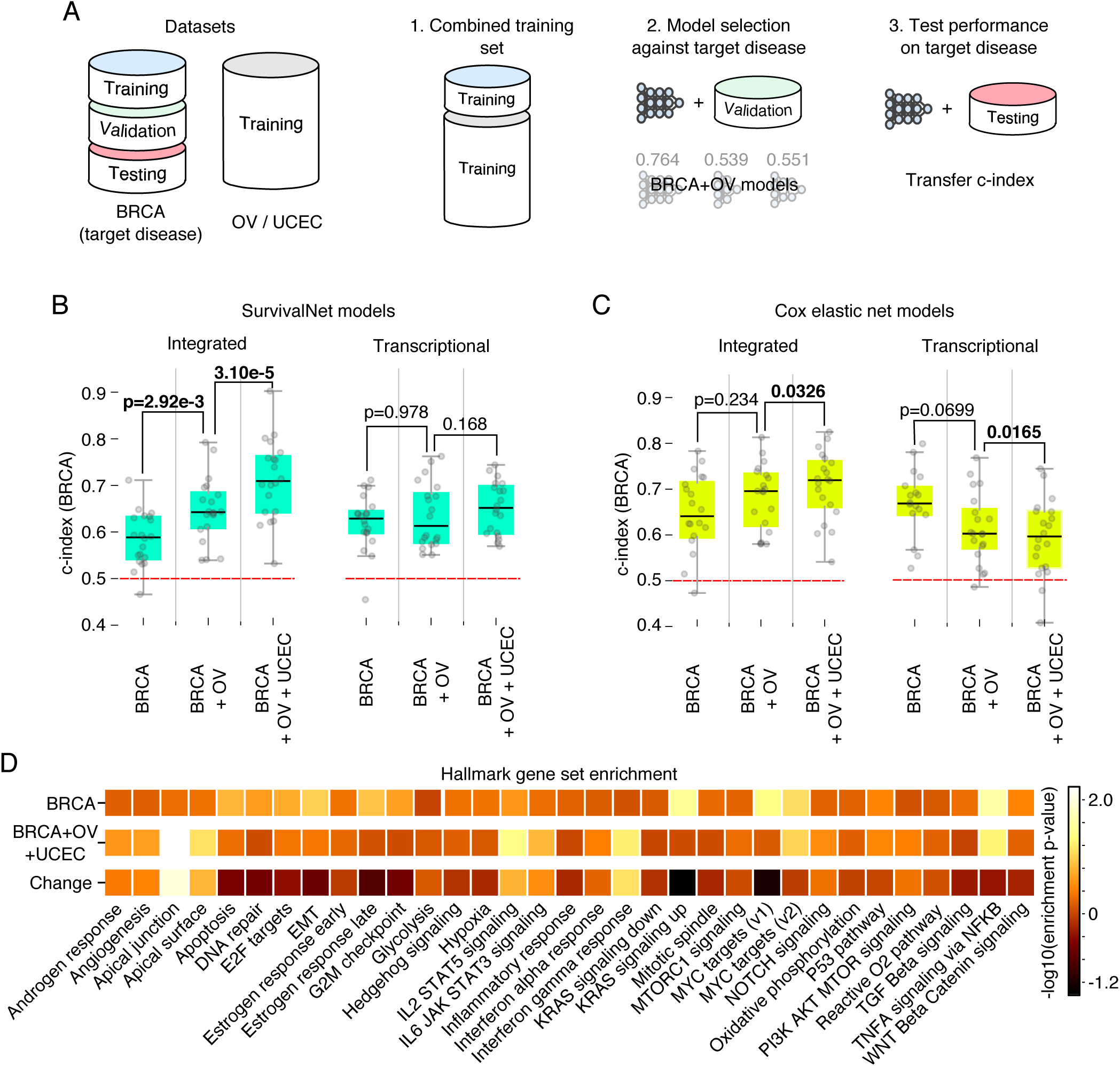
Learning with data from multiple cancer types improves deep survival models. **(A)** Data from the BRCA dataset was partitioned into training, validation, and testing sets. The BRCA training set was augmented with samples from the OV and UCEC and used to construct models for BRCA survival prediction. **(B)** Augmented training sets improve performance of SurvivalNet models for both the integrated and transcriptional feature sets. **(C)** For Cox elastic net, augmentation significantly degrades performance for the high-dimensional transcriptional feature set. **(D)** Gene set enrichment analysis of feature risk scores from the BRCA and BRCA+OV+UCEC transcriptional models. The model trained with BRCA+OV+UCEC samples emphasizes different biological concepts than the BRCA-only model.

Adding samples from the OV and UCEC datasets provides measurable improvements in BRCA prognostic accuracy for both integrated and transcriptional feature set deep survival models (see Figure 4B). For integrated models, training with BRCA+OV samples increases median c-index from 0.588 to 0.643 (rank-sum **p=2.92e-3**), and training with BRCA+OV+UCEC improves this further to 0.710 (**p=3.10e-5**). For the transcriptional feature set, training with BRCA+OV does not produce a measurable improvement over BRCA-alone (**p=0.978**), but training with BRCA+OV+UCEC provides a marginal improvement (**p=0.168**).

We also evaluated the ability of Cox elastic net to benefit from transfer learning, and found significant performance degradation with transfer learning in transcriptional feature set. Training with BRCA+OV reduces the median c-index to from 0.664 to 0.599 (**p=0.0699**), and training with BRCA+OV+UCEC reduces this further to 0.59335 (**p=0.0165**). Performance improvements with the integrated feature set for CEN were similar to those observed with deep survival models.

### Risk backpropagation analysis of transfer learning

To understand the information that OV and BRCA samples provide in predicting BRCA prognosis, we performed analysis of the BRCA and BRCA+OV+UCEC deep survival models using risk backpropagation. Risk backpropagation analysis was applied independently to the BRCA and BRCA+OV+UCEC transcriptional models to generate features risk scores, and gene set enrichment analyses were performed on these risk scores for each model to identify differences in pathway enrichment between the two models. Gene set enrichment scores for the BRCA+OV+UCEC model show increased emphasis on inflammatory pathways (particularly IL2-STAT5 signaling, IL6-JAK-STAT3 signaling and Interferon gamma response) as well as the apical junction gene set (known for its relevance to cell adhesion and metastasis). KRAS signaling and MYC targets v1 gene sets were de-emphasized in the BRCA+OV+UCEC model, pointing to a less prominent role of these pathways in determining breast cancer disease progression (See Figure 5D and Supplementary Tables S5 and S6).

## DISCUSSION

We created a software framework for Bayesian optimization and interpretation of deep survival models, and evaluate the ability of optimized models to learn from high-dimensional and multi-cancer datasets. Our software enables investigators to efficiently construct deep survival models for their own applications without the need for expensive hand tuning of design hyperparameters, a process that is time consuming and that requires considerable technical expertise. We also provide methods for model interpretation, using the backpropagation of risk to assess the prognostic significance of features and to gain insights into disease biology. Our analysis shows the ability of deep learning to extract important prognostic features from high-dimensional genomic data, and to effectively leverage multi-cancer datasets to improve prognostication. It also reveals limitations in deep learning for survival analysis and the value of complex and deeply layered survival models that need to be further investigated.

SN models have slightly better prognostic accuracy on two of three learning tasks using 17,000+ transcriptional features (GBMLGG and KIPAN), where CEN performed better using the lower-dimensional 300-400 integrated features. The high dimensionality of the transcriptional feature set presents a more challenging prediction problem where algorithms are more likely to overfit training data noise. CEN models are regularized linear models that use data-driven feature selection to identify a core subset of informative features for linear prediction. Their linearity does not appear to limit performance in our experiments, as their accuracy is similar to deep learning models and surpasses RSF models. While the deep models can effectively learn survival from high-dimensional data, the feature-learning capabilities of layered nonlinear transformations did not translate into significant gains as has been demonstrated in general image classification or language processing tasks (10). Larger datasets may be needed to overcome overfitting issues and to reveal anticipated performance benefits of deep learning. Deep learning methods typically require large amounts of training data to effectively learn their many parameters (27), although empirical results in some applications have demonstrated otherwise (28). In our experiments data requirements were exacerbated by the need to allocate validation samples for hyperparameter optimization. Smaller testing sets also introduced considerable variance in performance measurements.

Risk backpropagation analysis of gliomas demonstrated that deep learning could discover key features in high-dimensional data, identifying important genetic alterations that define clinical molecular subtypes of gliomas. Survival of patients diagnosed with infiltrating glioma depends largely on age, histologic grade and classification into three molecular subtypes defined by mutations in the Krebs cycle enzyme *isocitrate dehydrogenase* (*IDH1* / *IDH2)* and co-deletion of chromosome arms 1p and 19q (1): 1. Wild-type IDH (IDHwt) gliomas have an expected survival of 18 months, and are overwhelmingly diagnosed as advanced grade IV glioblastoma 2. IDH mutant gliomas with co-deletion of 1p and 19q (IDHmut-codel) have the best outcomes, with some patients surviving 10 years or more and 3. IDH mutant gliomas that lack co-deletions (IDHmut-non-codel) have intermediate outcomes. Our network identified *IDH1* and *IDH2* mutations (ranks 9, 10) as strongly associated with better prognosis, consistent with the role of these mutations as the primary feature in glioma classification. While our network did not explicitly identify 1p and 19q deletions as strongly associated with better prognosis (ranks 45, 233), it did identify *CIC* mutations which are a signature of the IDHmut-codel subtype (CIC mutations occur in more than 50% of IDHmut-codel gliomas), and *SMARCA4* mutations which occur in the less aggressive IDHmut-codel and IDHmut-non-codel subtypes. The top-ranked feature associated with poor prognosis in our analysis was deletion of *CDKN2A* deletion which is strongly associated with the aggressive IDHwt gliomas, as well as with a subset of poor prognosis IDH-mutated gliomas that lack broad DNA hypermethylation (GCIMP-low) (29). Loss of *PTEN* (rank 8) is also characteristic of aggressive IDHwt gliomas, has been shown to be an early event in gliomagenesis, and related to the loss of its parent chromosome 10 (10q and 10p were ranked 2 and 4, respectively) (30). Similarly, enrichment analysis of our transcriptional glioma model risk scores identified molecular pathways and processes related to epithelia to mesenchymal transition, a process that is associated with poor prognosis in cancers generally and specifically in gliomas.

Transfer learning experiments showed that deep survival models have an advantage in extracting prognostic information from multi-cancer datasets in the high-dimensional transcriptional feature set. Combining BRCA, OV and UCEC transcriptional data significantly degraded the accuracy of Cox elastic net models in predicting BRCA outcomes, but provided a small benefit to deep survival models. Augmentation of training datasets is a common technique in deep learning to compensate for small sample sizes. In this case, adding data from tumors with similar biology to the training set enhanced the performance of transcriptional and integrated models. Although genetic alterations and expression patterns are often strongly associated with primary disease site, common mechanisms of progression are likely shared by many cancers, and deep survival models can benefit from training with augmented datasets that provide additional evidence of these mechanisms. Enrichment analysis of risk scores from the BRCA-only and BRCA+OV+UCEC transcriptional models showed changes in the biological themes associated with highly prognostic transcripts, with increased emphasis on inflammatory response and cell adhesion in the BRCA+OV+UCEC model.

Although our study provides important insights into the use of deep learning for survival modeling, it has some limitations. Larger genomic datasets with clinical follow-up are needed to determine if the feature learning and nonlinearity of deep learning methods can provide substantial benefits in predicting survival. Secondly, our risk backpropagation analysis was simplified by averaging feature risk scores across patients. With nonlinear models, feature risk scores can vary significantly from patient to patient, and an in-depth analysis of these variations could yield insights into alternative paths for disease progression.

## METHODS

### Data

Al datasets were created using TCGAIntegrator (https://github.com/cooperlab/TCGAIntegrator), a Python module for assembling integrated TCGA genomic and clinical datasets with the Broad Institute Firehose (https://gdac.broadinstitute.org/). Datasets were filtered to remove patients lacking essential data platforms required in each experiment. Clinical variables including age and stage were required for each experiment, with missing radiation treatment status (binary) being mean-imputed to reflect prior likelihood in receiving radiation therapy. Features with categorical or ordinal values (i.e. stage) were expanded to a series of binary variables for model training. Copy number features were derived from the Affymetrix Genome-Wide Human SNP Array 6.0 platform. Gene expression features were taken as RSEM values from the Illumina HiSeq 2000 RNA Sequencing V2 platform. Protein expression measurements were taken from the MD Anderson Reverse Phase Protein Array (RPPA) Core platform that measures expression of cancer-relevant proteins and phosphoproteins. Sparse missing values in protein or gene expression features were 1nn-imputed (<20% missing values), where features exceeding this missing value threshold were discarded. Significant mutations were identified for inclusion in each dataset (LGG/GBM, KIPAN, BRCA) using a MutSig2CV <= 0.1 q-value threshold. Gene-level copy number features were filtered using a GISTIC <= 0.25 q-value threshold to identify focal events, and were further filtered using the Sanger Cancer Gene Census (31). All clinical and molecular features were standardized to zero-mean unit-variance to comply with best practices for training deep-learning algorithms. All datasets used to create this paper, along with the TCGAIntegrator commands used to generate these datasets are available on request.

### Software and hardware

All software used in training deep survival models, bayesian optimization, and model interpretation are provided as an installable python package at https://github.com/CancerDataScience/SurvivalNet. We have also provided a Docker container containing an installation of the package and all dependencies that provides access to SurvivalNet functionality without the need for software installations. SurvivalNet is implemented on top of the Numpy (v1.11) / SciPy (v0.18) stack using Theano (v0.8.2). Bayesian optimization was performed using the BayesOpt package (https://github.com/rmcantin/bayesopt). Survival analysis statistics like Kaplan Meier analysis and logrank testing were performed using the Python lifelines package (v0.8.0.1). Cox elastic net models were trained using Glmnet for Matlab (http://web.stanford.edu/∼hastie/glmnet_matlab/). Random survival forest models were trained using the RandomForestSRC (2.2.0) R package. Experiments were performed on a workstation equipped with two Intel Xeon E5-2620 v3 six-core processors, 64GB RAM, and two Titan-X GTX graphics processing units.

### Training, model selection and validation procedures

Deep survival models are multi-layer feed forward artificial neural networks with a Cox proportional hazards output layer that calculates negative log partial likelihood

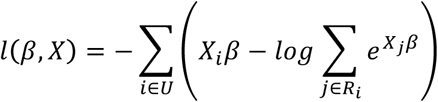

where *X*_*i*_ are the inputs to the output layer, *β* are the Cox model parameters, *U* is the set of right-censored samples and *R*_*i*_ is the set of “at-risk” samples with survival or follow-up times *Y*_*j*_ ≥ *Y*_*i*_.

This likelihood was optimized using backpropagation and line-search gradient descent. In each backpropagation iteration, the log partial likelihood is backpropagated throughout the network layers to update the interconnecting weights. The derivative used in backpropagation is

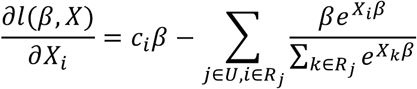

where *X*_*i*_ is the input to the output/Cox layer. This derivative is multiplied by derivatives of the hidden layers using the chain rule to update all the network parameters back to the first network layer. Training was performed in a single-batch configuration since likelihood calculations are not sample-independent. The network was regularized using random dropout of network weights.

Bayesian optimization was performed by splitting samples into training (60%), validation (20%) and testing (20%) sets. The training and validation sets were used by Bayesian optimization to determine the optimal model hyperparameters, namely number of layers (1-5), layer width (10-1000), dropout fraction (0-0.9) and activation function (Rectified-linear or hyperbolic tangent). The optimal model architecture was then applied to the testing set to evaluate c-index of the selected model. We repeated this procedure on 20 randomized assignments of the samples to training/validation/testing and performed 20 randomizations for each experiment.

Cox elastic net models contain two hyparparameters, *λ* which controls the overall degree of regularization and the mixture coefficient α that controls the balance between L2 and L1 norm penalties. Grid search over λ, α was performed to optimize the choice of these parameters. For each choice of α, a separate *λ* sequence was generated by Glmnet since the range of *λ* depends strongly on the α. A model was trained for each α / *λ* pair using the training set, and the model with the best performance on the validation set was then evaluated on the testing set. The same validation procedure was used to tune RSF hyperparameters including the number of trees (50, 100, 500, 1000), node size (1, 3, 5, 7, 9), and random splitting based on the recommendations in the randomForestSRC R package.

### Risk backpropagation and model interpretation

The models used for risk backpropagation and interpretation were created by identifying the best performing model configuration from the 20 randomized experiments. These configurations were then used to re-train a model using all available samples. Risk backpropagation was implemented using Theano to calculate the partial derivatives of risk with respect to each input variable using the multivariable chain rule. Given deep survival model with *H* hidden layers that operates on an *N*-dimensional feature vector *f* to predict risk *R*, the feature risk scores are calculated as the partial derivative of the model with respect to inputs

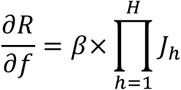

Where *J*_*h*_ is the Jacobian matrix of the *h*-th hidden layer with respect to its inputs, and *β* is the vector of parameters of the final layer that is a linear transformation (note the exponential is not applied since we are dealing with risk). This partial derivative is evaluated using the features of each patient *f*_*i*_ to generate an *N*-dimensional feature risk score vector for each patient. Features were ranked by calculating the median risk score for each feature across all patients.

For transcriptional models, feature risk scores were analyzed using the Preranked Gene Set Enrichment Analysis (GSEAPrerankedv1) module in GenePattern. The Hallmark gene set (32) from the MSigDB database (http://software.broadinstitute.org/gsea/msigdb/) was used for enrichment analysis. The HUGO Gene Nomenclature Committee database was used to harmonize gene symbols between gene sets and model features prior to GSEA analysis (http://www.genenames.org/).

### Transfer learning experiments

Datasets were combined using their shared features. For transcriptional and molecular features this merging is trivial, although many of the mutations and copy-number variations are dataset specific since they are filtered by GISTIC and MutSig to identify frequent alterations for each disease (integrated feature sets used in transfer learning are considerably smaller as a result). Pathologic stage and clinical stage were merged as a single “stage” variable where necessary, since their definitions of stage are similar (although the method of determining this stage differs). No additional normalization measures were employed to remove disease-specific biases.

### Data availability

This paper was produced using large volumes of publicly available genomic data. The authors have made every effort to make available links to these resources as well as making publicly available the software methods used to produce the datasets, analyses, and summary information. All data not published in the tables and supplements of this article are available from the corresponding author on request.

## AUTHOR CONTRIBUTIONS

SY and FA developed the primary method. SY, FA curated datasets. SY, FA, CD, JL, DAG performed experiments. CS assisted with software development. MA, SH, JEVV, and DJB assisted with interpretation of experimental results. LAD conceived of the concepts and oversaw the work.

## ACKNOWLEDGEMENTS

This work was supported by the National Brain Tumor Society Oligo Research Fund, U.S. National Institutes of Health, National Library of Medicine Career Development Award K22LM011576, and National Cancer Institute grant U24CA194362, National Institutes of Health CTSI grants UL1TR000454 and TL1TR000456, and with funds from the Emory Winship Cancer Institute.

